# PSX: Protein-Solvent Exchange - Software for calculation of deuterium-exchange effects in SANS measurements from protein coordinates

**DOI:** 10.1101/609859

**Authors:** Martin Cramer Pedersen, Yong Wang, Frederik Grønbæk Tidemand, Anne Martel, Kresten Lindorff-Larsen, Lise Arleth

## Abstract

Recent developments in neutron scattering instrumentation and sample handling have enabled studies of more complex biological samples and measurements at shorter exposure times. The experiments are typically conducted in D_2_O-based buffers to emphasize or diminish scattering from a particular components or to minimize background noise in the experiment. To extract most information from such experiments it is thus desirable to determine accurate estimates of how and when closely bound hydrogen atoms from the biomolecule exchange with the deuterium in the solvent. We introduce and document software, PSX, for exploring the effect of hydrogen-deuterium exchange for proteins solubilized in D_2_O as well as the underlying bioinformatical models. The software aims to be generally applicable for any atomistic structure of a protein and its surrounding environment, and thus captures effects of both heterogenous exchange rates throughout the protein structure and by varying the experimental conditions such as *pH* and temperature. The paper concludes with examples of applications and estimates of the effect in typical scenarios emerging in small-angle neutron scattering on biological macromolecules in solution. Our analysis suggests that the common assumption of 90% exchange is in many cases an overestimate with the rapid sample handling systems currently available, which leads to fitting and calibration issues when analysing the data. Source code for the presented software is available from an online repository in which it is published under version 3 of the GNU publishing license.

## 1. Introduction

Small-angle neutron scattering (SANS) experiments are a powerful source of information on the structure and dynamical properties of macromolecules such as proteins. In order to extract as much information as possible from such experiments during, for example, model refinement, it is often necessary to provide external information such as scattering lengths, molecular volumes, and similar physical parameters. SANS experiments are often carried out using either D_2_O or H_2_O/D_2_O mixtures providing opportunities for varying contrast between different parts of the sample, due to the stark difference in scattering properties between protons and deuterons. When a protonated protein is exposed to D_2_O, a number of protons in the protein will exchange with the deuterons in the solvent over a timescale that may be comparable to that of the experiment. Thus, for biomolecular samples and experiments relying on D_2_O-based buffers (such as SANS and neutron reflectometry), this exchange of protons must be accounted for in order to refine accurate models of the samples in question. These phys-ical effects have been discussed for decades (Stuhrmann, 1974; Stuhrmann, 1978; Zaccai & Jacrot, 1983) in the bio-scattering community and in particular in the context of contrast variation experiments in which a sample is irradiated by neutrons in a series of buffers with varying D_2_O content to enhance the contrast of different parts of the sample (Stuhrmann, 1974; Whitten *et al*., 2007; Clifton *et al*., 2013). The paper by Stuhrmann (Stuhrmann, 1974) explicitly measures the H-D exchange rate for myoglobin by considering the change in zero-angle scattering over time. Stuhrmann observes a rapid exchange happening on the order of a second, and a slower exchange occuring on the order of minutes.

These considerations become increasingly relevant for proteins with domains shielded from the solvent such as e.g. a membrane protein in a phospholipid bilayer nanodisc (Wadsäter *et al*., 2012; Kynde *et al*., 2014; Johansen *et al*., 2018), liposomes (Rubinson *et al*., 2013), cubic lipid-phase (van’t Hag *et al*., 2016), detergent micelle (Midtgaard *et al*., 2018), or proteins complexed with e.g. other proteins (Whitten *et al*., 2007; Clifton *et al*., 2012) or RNA (Gabel, 2015). The recent developments of so-called “invisible” carrier systems that match the scattering properties of the associated solvent, have been used for solubilizing membrane proteins for solution SANS experiments; in particular the so-called match-out deuterated detergent micelles (Midtgaard *et al*., 2018) or stealth nanodiscs (Maric *et al*., 2014; Josts *et al*., 2018), add further relevance to the considerations.

On a similar note, in-line size exclusion chromatography (SEC) for BioSANS and stopped-flow time-resolved SANS (Rössle *et al*., 2000) have recently emerged as a promising technique for investigating particularly complicated and fragile biological samples of interest (Jordan *et al*., 2016; Johansen *et al*., 2018). When employing this technique, samples are exposed to the solvent, in many cases D_2_O, only minutes or even seconds before being irradiated by the neutron beam, meaning that the exchange pro-cess is far from equilibrium. Conversely, in traditional SANS experiments for weakly scattering samples can be irradiated for days or even weeks. Equally relevant are the sample preparation protocols, in which some samples spent days in D_2_O before irradiation.

In this paper, we present an approach to and software for more accurately estimating the hydrogen-to-deuterium exchange (also commonly known as protium-deuterium exchange) in a protein structure as a function of the time in which the protein has been exposed to D_2_O using a well established model proposed by Vendruscolo and colleagues (Vendruscolo *et al*., 2003; Best & Vendruscolo, 2006) from the world of structural bioinformatics. The software focuses on the exchange of the backbone hydrogens, as these are the most relevant exchanges in the context of neutron scattering experiments (Figure 1). Specifically, we divide hydrogens into three groups: (1) non-exchanging, carbon-bound hydrogens, (2) slowly-exchanging backbone hydrogens that exchange with the solvent at timescales relevant for neutron scattering experiments on biological samples, which exchanges with the solvent in timescales of minutes, several days or even longer, and (3) the remaining rapidly-exchanging side chain hydrogens, which exchange with the solvent at subsecond timescales.

**Fig. 1.**
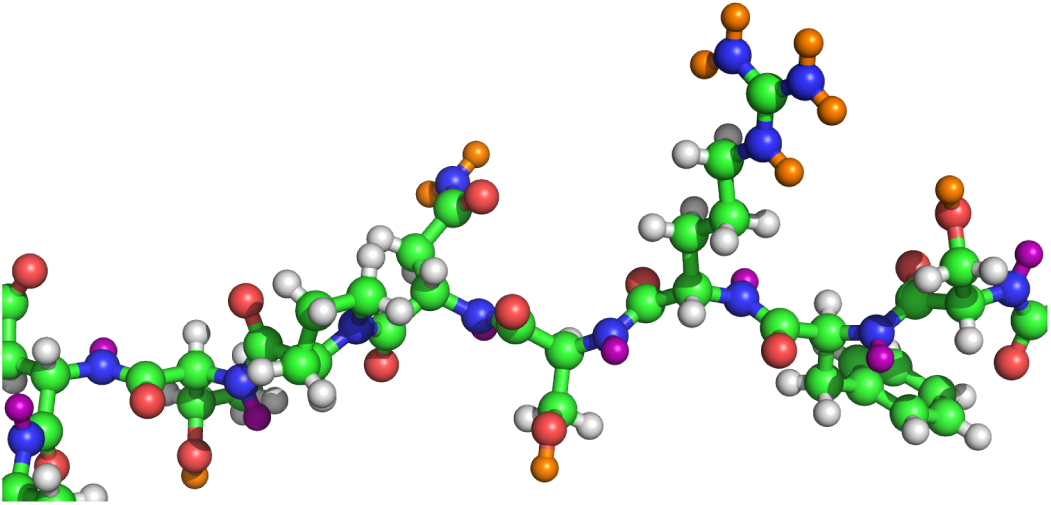
A generic protein structure with backbone amide hydrogens highlighted in purple, un-exchangeable hydrogens in white, and the rapidly-exchanging hydrogens in orange. Carbon-, nitrogen-, and oxygen-atoms are shown in green, blue, and red, respectively.

The hydrogen-deuterium exchange phenomenon is routinely observed and investigated in neutron protein crystallography (Blum *et al*., 2007; Blakeley *et al*., 2008; Blum *et al*., 2009), where samples are often either grown in or soaked in D_2_O-based buffers before irradiation, as the technique is particularly sensitive the differences between hydrogen and deuterium and the presence of one isotope is often preferred in specific parts of the sample. Furthermore, partial exchange of the backbone amino acid hydrogens is reported; in line with the model introduced in the previous paragraph. Using this technique, back-exchange for deuterated crystals in hydrogen-based solvents is observed as well (Yee *et al*., 2017).

Some existing software packages such as CRYSON (Svergun *et al*., 1998) and ISIS’ Biomolecular Scattering Length Density Calculator (ISIS, 2019) assume a homogeneous deuteration level throughout the molecule (defaulting to 90% of the backbone hydrogens and 100% of the rapidly-exchanging hydrogens in CRYSON and 90% of the rapidly exchanging hydrogens plus the backbone hydrogens in ISIS’ Biomolecular Scattering Length Density Calculator); thus not accounting for actual solvent exposure and the geometry of the protein. Other software packages such as e.g. SASView (The SASView Project, 2019) or WillItFit (Pedersen *et al*., 2013) implicitly rely on accurate estimates of scattering lengths. The presented software will aid users in preparing their models for these software packages and for estimating the effects of the exchange effect for their samples.

## 2. Empirical model for hydrogen-deuterium exchange

We here describe an approach to estimate the level of deuteration of each of the exchangeable backbone amide protons using the structure of the protein. We do so via estimating both the intrinsic exchange rate constants from the amino acid sequence, i.e. the exchange rate for an exposed, entirely unfolded amino acid residue, and the perturbation away from these intrinsic rates in a given conformation of the protein. We assume that hydrogen-deuterium exchange can be described by the well-established Linderstrøm-Lang model (Hvidt & Nielsen, 1966), in which the exchange of a buried or otherwise protected proton with a solvent deuteron processes in two steps. We here only provide a brief description of this model, and refer the reader to the literature for a more in-depth discussion (Hvidt & Nielsen, 1966; Englander & Kallenbach, 1983; Teilum *et al*., 2005).

In the Linderstrøm-Lang model each site is treated as existing either in a “protected” (closed) state or an “exchange competent” state (open) state. In order for exchange to occur, the site thus needs to become transiently exposed to the solvent going from the closed to the open state in a process with an equilibrium constant termed *K*_open_. Under certain experimental condition the rate of opening and closing affects the overall exchange rate, but under the conditions and assumptions explored here, only the equilibrium constant is important. During this transient exposure, the site may exchange with the solvent with an intrinsic rate constant, *k*_int_, determined by the local sequence and the experimental conditions. The intrinsic rate is the rate at which a fully exposed amide would exchange under the given set of conditions. Solving the rate equations for this model leads to a two-exponential exchange process, but after a rapid transient that is generally over in less than a second (Teilum *et al*., 2005), the exchange process can be described by a pseudo-first order reaction with a rate constant, *k*_ex_. In this way, the probability, *P*_ex_, of a backbone hydrogen having exchanged with a hydrogen atom from the solvent becomes:

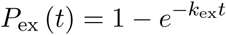

where *t* is the time elapsed since the exposure to D_2_O. Because amides are, to various extents, protected from exchange by burial and hydrogen bonding, *k*_ex_ *≤ k*_int_.

Under relatively general assumptions and experimental conditions (leading to the so-called EX2 regime (Hvidt & Nielsen, 1966; Teilum *et al*., 2005)), the observed exchange rate, *k*_ex_, is simply the product of the equilibrium constant for transient opening, *K*_open_, and the intrinsic rate, *k*_int_. Briefly explained, this occurs when the rate of “closing” (i.e. going from the exchange competent state back to the protected state) is substantially faster than the intrinsic rate (Teilum *et al*., 2005). We note here that not all amides exchange with EX2 kinetics, though such exchange kinetics is often found at pH values up to an somewhat above neutral pH. At higher pH, where *k*_int_ is often very large one instead often finds the so-called EX1 regime where the rate of opening (leading to the exchange competent state) limits exchange kinetics (Teilum *et al*., 2005).

Mathematically, the exchange rate in the EX2 regime is typically formulated through the so-called protection factor, 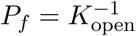:

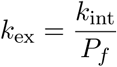

Thus, summarized the overall exchange rate *k*_ex_ at a site is determined by combining the intrinsic exchange rate, *k*_int_, at that site and the level of protection afforded at the site (*P_f_*) by the surrounding protein structure through e.g. burial from solvent or involvement in hydrogen bonds.

The structural dependencies of the exchange rate are thus captured through differences in protection factors. Here we describe these using a model originally presented by Vendruscolo and colleagues (Vendruscolo *et al*., 2003) in which the level of protection is predicted from examining the number of contacts and hydrogen bonds formed at an exchanging site:

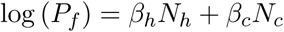

where *N_h_* is the number of hydrogen bonds to the backbone nitrogen/hydrogen, and *N_c_*is the number of heavy atoms within a distance of 6.5 Å of the backbone nitrogen. For atomistic models, the coefficients *β_h_* = 2.0 and *β_c_* = 0.35 were refined from an variety of molecular dynamics (MD) simulations and data from hydrogen exchange (HX) experiments (Best & Vendruscolo, 2006). In the following examples and discussions, we estimate the presence of hydrogen bonds using the DSSP algorithm (Kabsch & Sander, 1983) with a default energy threshold of *−*0.5 kcal/mol. We note that since this relationship was parameterised and validated from data on seven small proteins with HX data at near-ambient conditions, we expect that the accuracy of this model is limiting in our application and that it should not be applied to conditions very different from those used in these original studies (Best & Vendruscolo, 2006).

We show examples of protection factors predicted as described above on their respective atomic structures (Figure 2). As expected, we observe how well folded domains are more protected than flexible, solvent exposed regions. Neutron crystallography experiments qualitatively support these predictions (Blum *et al*., 2007; Blakeley *et al*., 2008; Blum *et al*., 2009). Furthermore, we note that transmembrane regions of membrane proteins are considered to be very well protected from the solvent; again in line with intuition. We note that in case of calculations for membrane proteins we embed the protein in a lipid bilayer or other membrane mimetic, and take contacts to these into account when calculating *P_f_*.

**Fig. 2.**
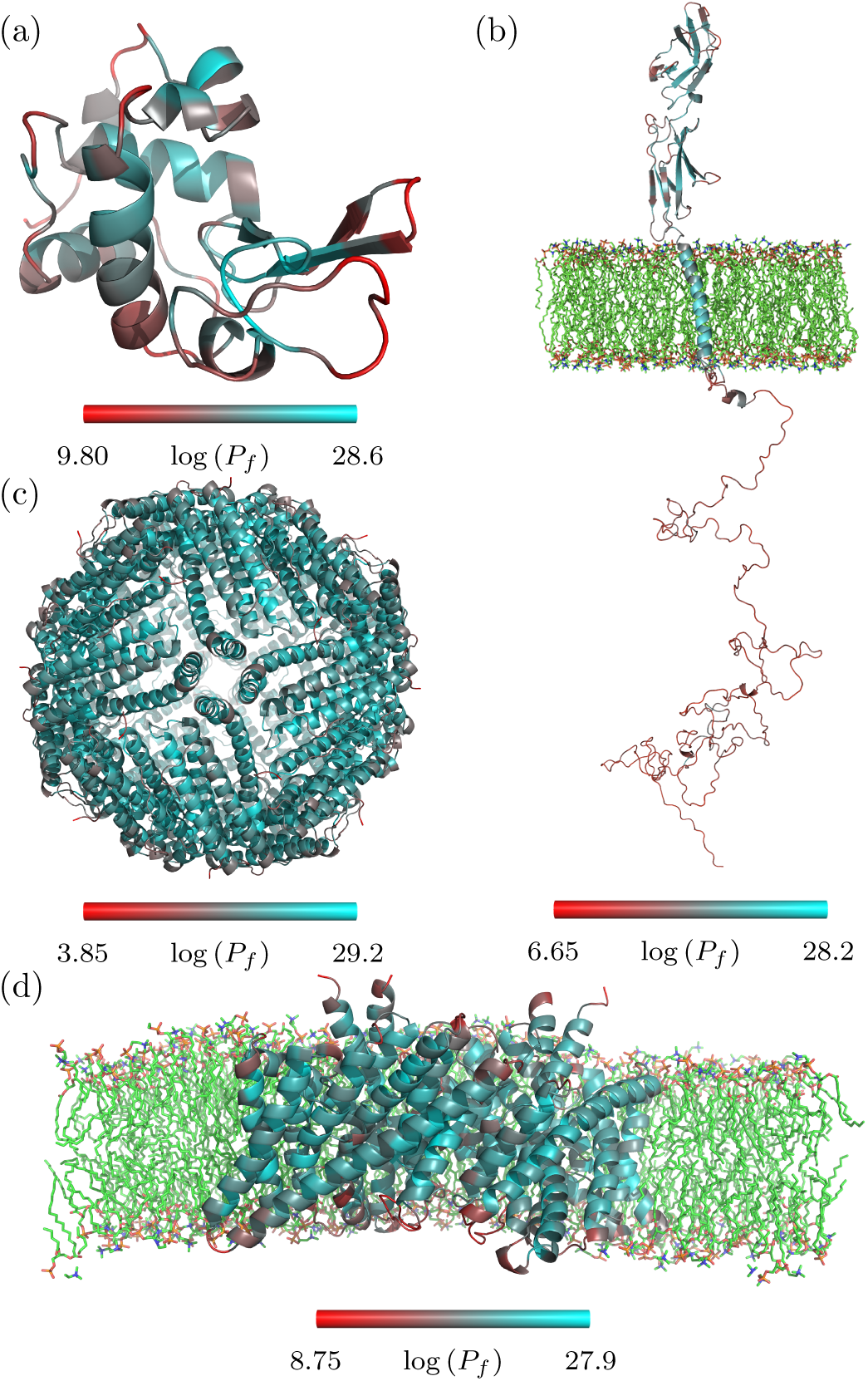
Examples of protection factors calculated from protein structures. (a): Lysozyme, from PDB ID **2LYZ**. (b): Prolactin receptor in a bilayer of POPC phospholipids (half of which are not shown), from Bugge *et al*. (Bugge *et al*., 2016). (c): Apoferritin (24-mer), from PDB ID **1IES** (Granier *et al*., 1997). (d): H^+^/Cl^−^ Exchange Transporter in a bilayer of DPPC phospholipids (half of which are not shown), from PDB ID **1OTU** (Dutzler *et al*., 2003).

The intrinsic exchange rate, *k*_int_, represents the exchange rate for a fully unfolded peptide chain. It is readily estimated from the amino acid residue sequence (and the experimental conditions) using e.g. the Sphere server (Zhang, 1995; Fox Chase Cancer Center, 2019), which bases its estimations on values found in the literature (Bai *et al*., 1993; Connelly *et al*., 1993) or the data published by the Englander Lab (Englander, 2019; Nguyen *et al*., 2018), which cites another source for the high-*pH* exchange rates (Mori *et al*., 1997). The software presented in this paper uses the values from the latter two sources, which are refined from HX experiments on various reference peptides.

In a Gedankenexperiment imagining a measured *pH* of 7 at a temperature of 293.15 K, we estimate the intrinsic exchange rates for the sequence of lysozyme range from s*^−^*^1^ to 46.60 s*^−^*^1^ across the 129 amino acid residues. For comparison, the same calculation at a temperature of 283.15 K yields intrinsic exchange rates from 0.01 s*^−^*^1^ to 16.62 s*^−^*^1^ for the same sequence. Similarly, with a measured *pH* of 8 at a temperature of 293.15 K, we obtain exchange rates from from 0.29 s*^−^*^1^ to 465.95 s*^−^*^1^ for our lysozyme.

Apart from the backbone hydrogens, we assume that the following hydrogen atoms will exchange with the solvent at a sub-second timescale due to their low *pK_a_* values (Lide, 2005); much faster than the ones relevant for neutron scattering experiments:

- The – NHs in the N-terminal(s)
- The – OHs in SER, THR, and TYR
- The – SHs in CYS
- The side chain – NHs in ASN, GLN, ARG, LYS, HIS, and TRP

In other words, these chemical groups are assumed to instantly exchange hydrogen atoms with the solvent. We refer to these atoms as rapidly-exchanging throughout this paper. The set of rapidly-exchanging hydrogens and backbone hydrogens are sometimes referred to as the “labile” hydrogens in the literature. We note that in some cases exchange of the single NH group in the indole group of TRP residues may be rather slow (Wedin *et al*., 1982), but since TRP residues are rather rare (approximately 1.5% of residues in folded proteins (Tompa, 2002)) and there is no quantitative model for its exchange, we consider them as generally rapidly exchanging. We acknowledge that this is an approximation that can be removed should a better model for intrisic rates and protection of indoles (or other non-NH groups) become available. The remaining hydrogens are considered non-exchangeable (Figure 1).

As an example of the result of these experimentally-supported assumptions, out of the 959 hydrogen atoms generated by PDB2PQR (Dolinsky *et al*., 2004; Dolinsky *et al*., 2007) for the PDB ID **2LYZ** (Diamond, 1974) structure of lysozyme consisting of 129 amino acid residues at a measured *pH* of 7.0, we find 148 rapidly-exchangeable, 126 backbone hydrogens (for a total of 274 labile hydrogens), and the remaining 685 non-exchangeable.

## 3. Solution small-angle neutron scattering

### 3.1. Estimating scattering from a protein in solution

For the calculations of the scattering profiles (i.e. the excess scattered intensity, *I*(*q*), as a function of the scattering momentum transfer, *q*) presented here, we have used a simple fast Debye sum (Hansen, 1990) based on commonly used scattering lengths (NIST, 2019) and volumes (Fraser *et al*., 1978) (Table 1). We assume a partial specific volume of solvent D_2_O of 30.0 Å^3^. Details of these calculations can be found in Appendix A.

**Table 1.** The (coherent) scattering lengths and volumes used in the calculation of scattering profiles for proteins presented in this paper. The values are from NIST (NIST, 2019) and the literature (Fraser et al., 1978), respectively.

| Atom | Scattering length, fm | Assigned volume, Å<sup>3</sup> |
| --- | --- | --- |
| H | −3.7406 | 5.15 |
| D | 6.671 | 5.15 |
| C | 6.646 | 16.44 |
| N | 9.36 | 2.49 |
| O | 5.803 | 9.13 |
| S | 2.847 | 19.86 |

We note that more advanced and involved methods for estimating scattering profiles from the atomic coordinates exist; e.g. expansion of scattering amplitudes on spherical harmonics as employed by the software CRYSON (Svergun *et al*., 1998) or Debye sums as employed by the software FoXS (Schneidman-Duhovny *et al*., 2010) (for x-ray scattering). The simple, fast Debye method we employ appears to be capable to reproduce the data in Figure 4.

**Fig. 3.**
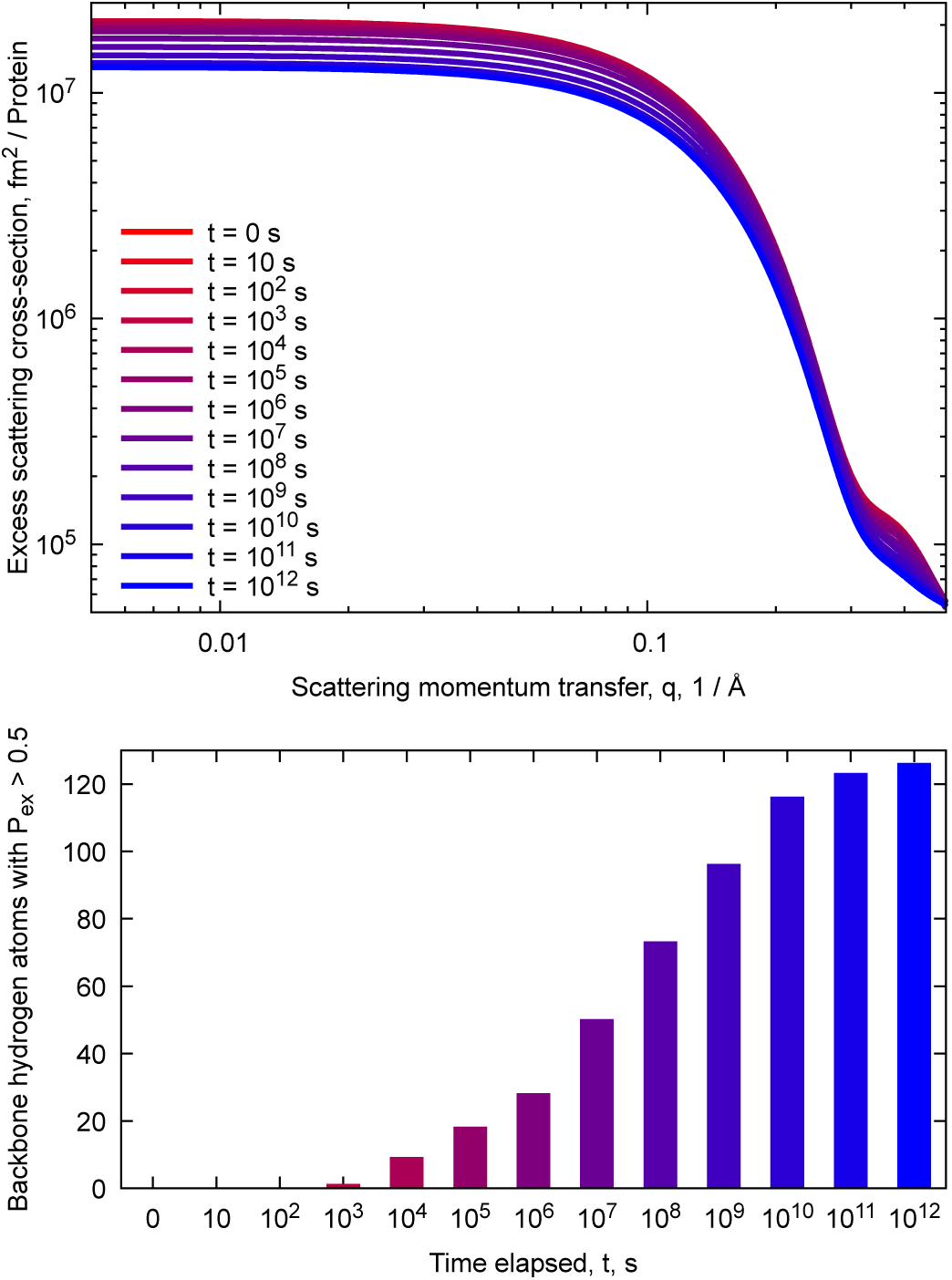
Time evolution of the structure PDB ID **2LYZ**, lysozyme, shown in Figure 2(a), according to the solvent exposure model discussed in this paper. Top: Scattering profiles calculated as explained in Section 3.1 and Appendix A; a small background has been added the the profiles for a more realistic presentation. Bottom: The number of the 126 backbone hydrogens atoms with *P*_ex_ *>* 0.5 for selected time points. We remind the reader that 10^5^ s is approximately 1.15 days and that 10^8^ s is approximately 3.16 years.

**Fig. 4.**
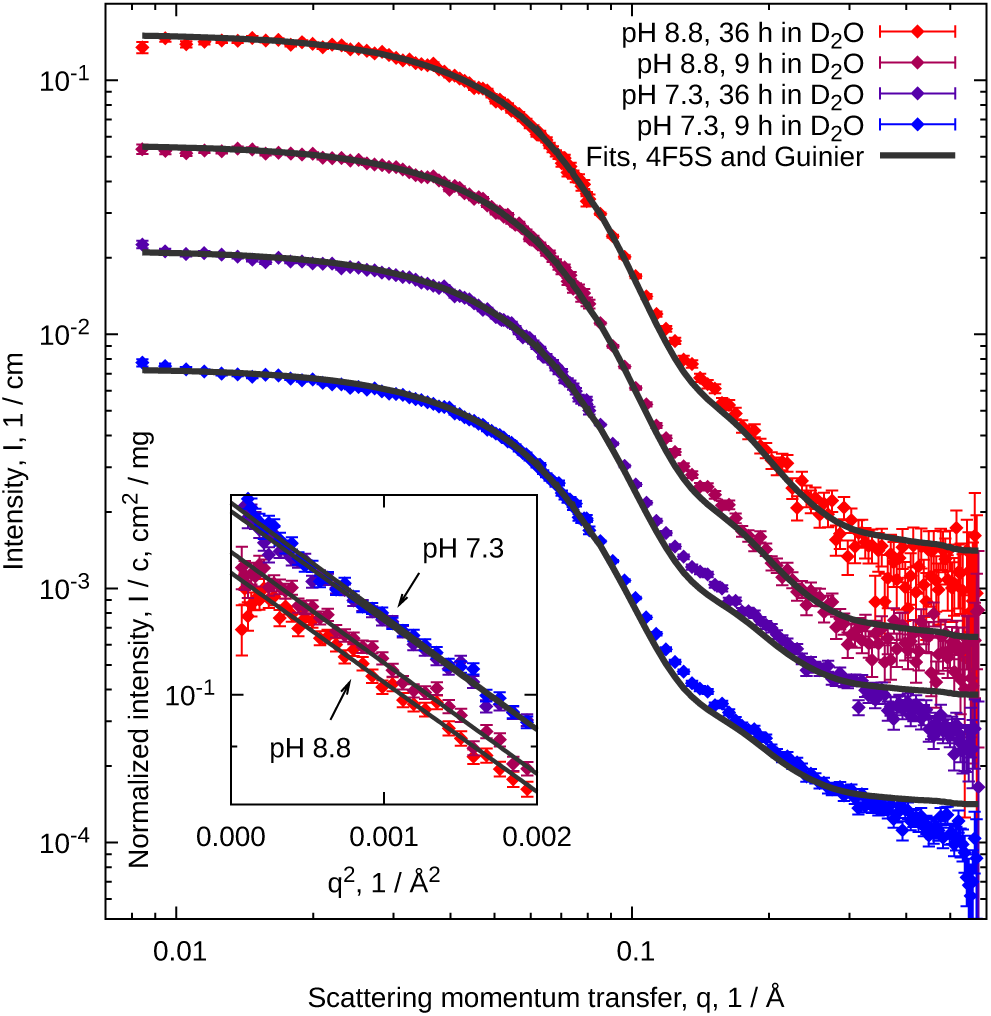
SANS data from BSA. The top-most data and fit are correctly scaled, whereas the lower data and fits are scaled by factors of 3 relative to the one above. The insert shows Guinier fits. Selected parameters are listed in Table 2.

**Table 2.** Concentration, c, radius of gyration, R_g_, and normalized forward scattering, I(q → 0)/c for the data presented in Figure 4. The Guinier fits (Guinier & Fournet, 1955) were done using Gnuplot (Williams et al., 2019).

| Sample | $c$, mg/ml | $R_g$, Å | $I(q \rightarrow 0)/c$, cm<sup>2</sup>/mg |
| --- | --- | --- | --- |
| ■ | 1.18 | $25.8 \pm 0.3$ | $0.1280 \pm 0.0007$ |
| ■ | 1.24 | $26.0 \pm 0.3$ | $0.1336 \pm 0.0006$ |
| ■ | 1.30 | $25.7 \pm 0.3$ | $0.1460 \pm 0.0007$ |
| ■ | 1.32 | $26.2 \pm 0.3$ | $0.1476 \pm 0.0007$ |

### 3.2. Comparison of time scales: Lysozyme

As a first example, let us consider the application of the presented scheme in a Gedanken experiment: a generic solution SANS experiment on lysozyme at room temperature, 293.15 K, in a D_2_O-based buffer with a measured *pH* of 7.0. In Figure 3, a calculation of the scattering profile at selected times is shown, along with the amount of backbone hydrogens expected to have exchanged with the solvent.

We observe that in order for the imagined lysozyme to obtain fully or even 90% deuterated backbone hydrogens, the relevant time scales are orders of magnitude beyond those relevant in an experimental experiment. Indeed, the model predicts that the 90% backbone hydrogen deuteration commonly assumed, would not be reached until between 10^9^ s and 10^10^ s or a timespan of more than 30 years under the defined conditions. As expected, the excess scattering cross-section for vanishing values of *q*, usually dubbed “the forward scattering”, decreases as the protein deuterates; as the exchange process brings the scattering length density of the protein closer to that of D_2_O. In total, the forward scattering drops to 62% of the initial value; see Figure 3 (top). It is particularly noteworthy that within typical experiment times of 1 hour and 1 day, the forward scattering intensity drops to, respectively, 98% and 94% of the initial value.

Regarding the shape of the curve, we observe a small change in the radius of gyration of the protein (data not shown) but interestingly note that the high-*q* behaviour changes during the exchange (Figure 3).

In particular, we note that of the 274 labile sites, only 54% are rapidly-exchanged, and after more than a day (10^5^ s) of exposure to D_2_O the model by Vendruscolo *et al*. predicts that only a total of 61% of the 274 labile hydrogen atoms will have exchanged. This value should be considered in relation to the usually applied 90% discussed earlier. We note also many residues in lysozyme have experimentally-determined values of *k*_ex_ *<* 10*^−^*^7^s*^−^*^1^ at *pH* 7.0 (Pedersen *et al*., 1993). This is fully consistent with the very long exchange times obtained from the model calculations.

For comparison, the effect of including a hydration layer using e.g. CRYSON (Svergun *et al*., 1998) decreases the scattering intensity by up to a few percent using default parameters making the exchange effect relatively more impactful.

### 3.3. Comparison to experimental data: BSA

The data presented in Figure 4 were collected at the SANS instrument D22 (D22, 2019), Institut Laue-Langevin, in Grenoble, France. Four samples of bovine serum albumin (BSA) were prepared from the same H_2_O-based stock solution at either *pH* 7.3 or *pH* 8.8 measured by *pH*-meter and changed into two analogous D_2_O-based buffers using a NAP-5 desalting column (GE healthcare) either 9 or 36 hours before irradiation. The experiment, including sample storage, was conducted at approximately 6 *^◦^*C. For further details on the sample preparation and the experimental set up, consult the appendix.

We used `PROPKA` (Olsson *et al*., 2011) and chain A of PDB ID **4F5S** (Bujacz, 2012) to estimate 4564 hydrogens in the BSA monomer, at *pH* 7.3 (*pD* of 7.7) and 4560 at *pH* 8.8 (*pD* of 9.2). According to the discussed model for H/D-exchange, the structure contains 499 hydrogen sites which will exchange rapidly (at *pH* 7.3 and an additional 4 at *pH* 8.8) with the D_2_O-based solvent and 554 slowly exchanging backbone hydrogens, the exchange rates of which we estimate using the presented software.

From Figure 4, we conclude that quantitively we observe the expected behaviour. We see the forward scattering, *I*(*q →* 0), decreasing as a function of how long time the proteins have been exposed to D_2_O (Figure 4, insert). Similarly, we note that the high-*pH* samples appear to have considerably lower forward scattering than the low-*pH* samples, in line with the expectation of more rapid exchange in these samples (Table 2). We note also that the effect of varying the time is greater at *pH* 8.8 where exchange is faster than at *pH* 7.3.

To fit the data we had to fit a scaling parameter which refined to 1.05 *−* 1.14 along with a small constant background for the four presented samples. While it is obviously unsatisfactory to have to fit these adjustment parameters, we consider the refined values to be well within the margin-of-error of the absolute calibration of our data.

### 3.4. Comparison of homogeneous and site-specific deuteration: SERCA

Let us consider the membrane protein sarco-/endoplasmatic reticulum Ca^2+^ ATPase (SERCA) in an equilibrium between the E1 state, from PDB ID **5XA7** (Norimatsu *et al*., 2017), and the E2 state, from PDB ID **5XAB** (Norimatsu *et al*., 2017), described by an ensemble of structures in a carrier system, which is assumed to be ‘invisible’ in D_2_O as previously discussed (Midtgaard *et al*., 2018; Maric *et al*., 2014; Josts *et al*., 2018).

We generated the ensemble of E1 and E2 states of SERCA using two independent one-microsecond MD simulations with CHARMM36m force field (Huang *et al*., 2016) initiating from the aforementioned crystal structures inserted into a 1-palmitoyl-2-oleoylphosphatidylcholine (POPC) membrane bilayer. The simulations used the TIP3P water model and a periodic box of 13 nm by 13 nm by 16 nm and were performed using Gromacs2016.3 (Abraham *et al*., 2015). In total, we generate an ensemble of 40 structures describing the intermediate states between the aforementioned crystal structures. As previously described we average the calculated protection factors over these structures (Best & Vendruscolo, 2006):

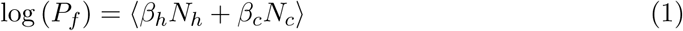

and show calculated scattering profiles for the described structure under various conditions (Figure 5) at a measured *pH* of 7.0 and a temperature of 293.15 K.

**Fig. 5.**
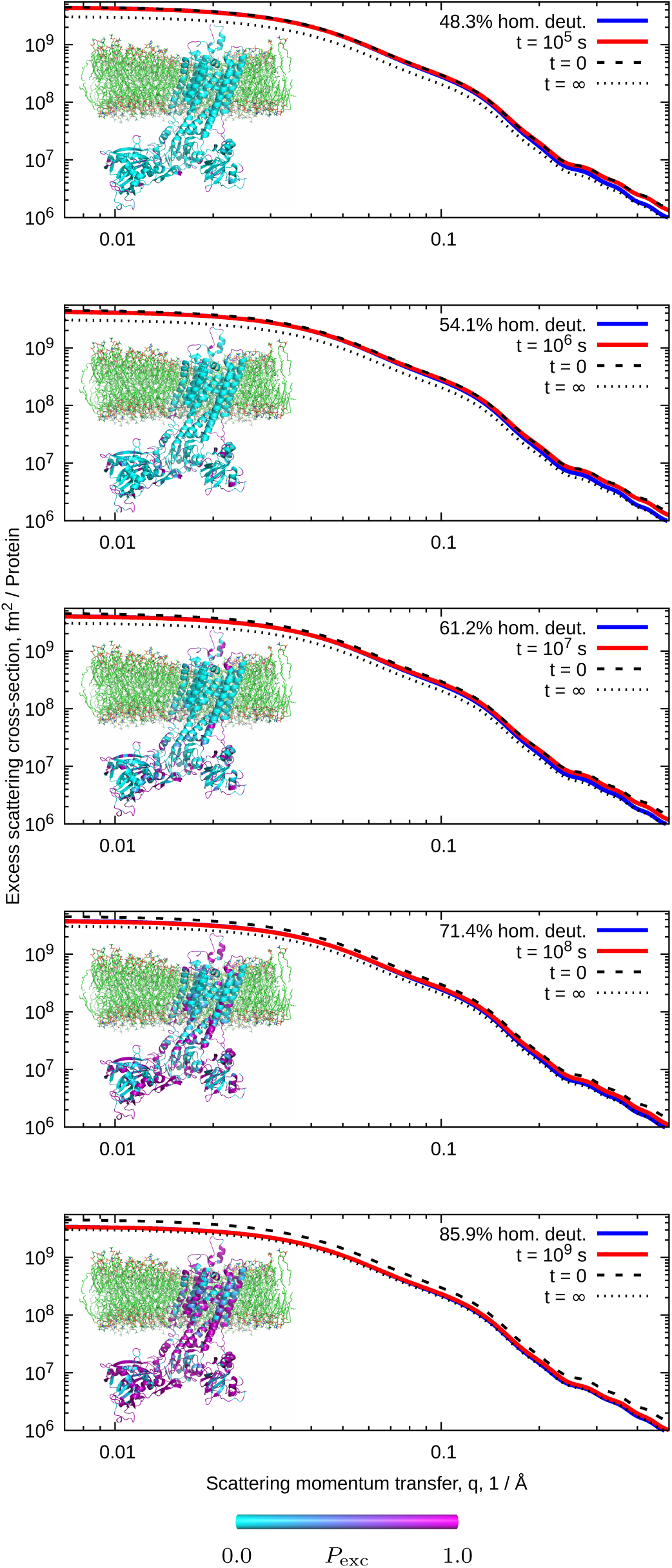
Scattering profiles of SERCA. As inserts, we visualize *P*_ex_ on the E1 state of SERCA in a bilayer of POPC phospholipids (half of which are not shown). The phospholipids do not contribute to the scattering pattern in an effort to mimick the scattering from match-out deuterated carrier systems. For reference, we also plot the scattering profiles of the fully deuterated structure and the structure with only the rapidly-exchanging sites exchanged to deuterium.

We observe how the model predicts that the extra-cellular domains of the protein exchange faster than the transmembrane parts buried in the bilayer and how the effect of specifically exchanging specific sites mostly impacts the shape of the scattering profile for *q >* 0.1 Å*^−^*^1^. In line with intuition, we observe that the difference between homogeneous deuteration and specific exchange matters most when the solvent-exposed domains of the proteins are mostly exchanged, while the transmembrane domains are still mostly protonated; i.e. the situation, wherein the protein appears mostly like a two-contrast molecule.

## 4. Software

### 4.1. Input and output

As input, the software takes the atomic coordinates in PDB-format (Bernstein *et al*., 1977) of a protein alongside the coordinates of any other atoms relevant to the calculation e.g. a lipid bilayer for a membrane protein or bound ligands. We emphasize that unique model and chain identifiers specifying the individual models and molecules must be specified in the file as the calculation of *k*_int_ depends on the chain structure of the protein. Several additional, optional arguments can be supplied for the calculation. A list of the most important optional arguments are listed in Table 3.

**Table 3.**
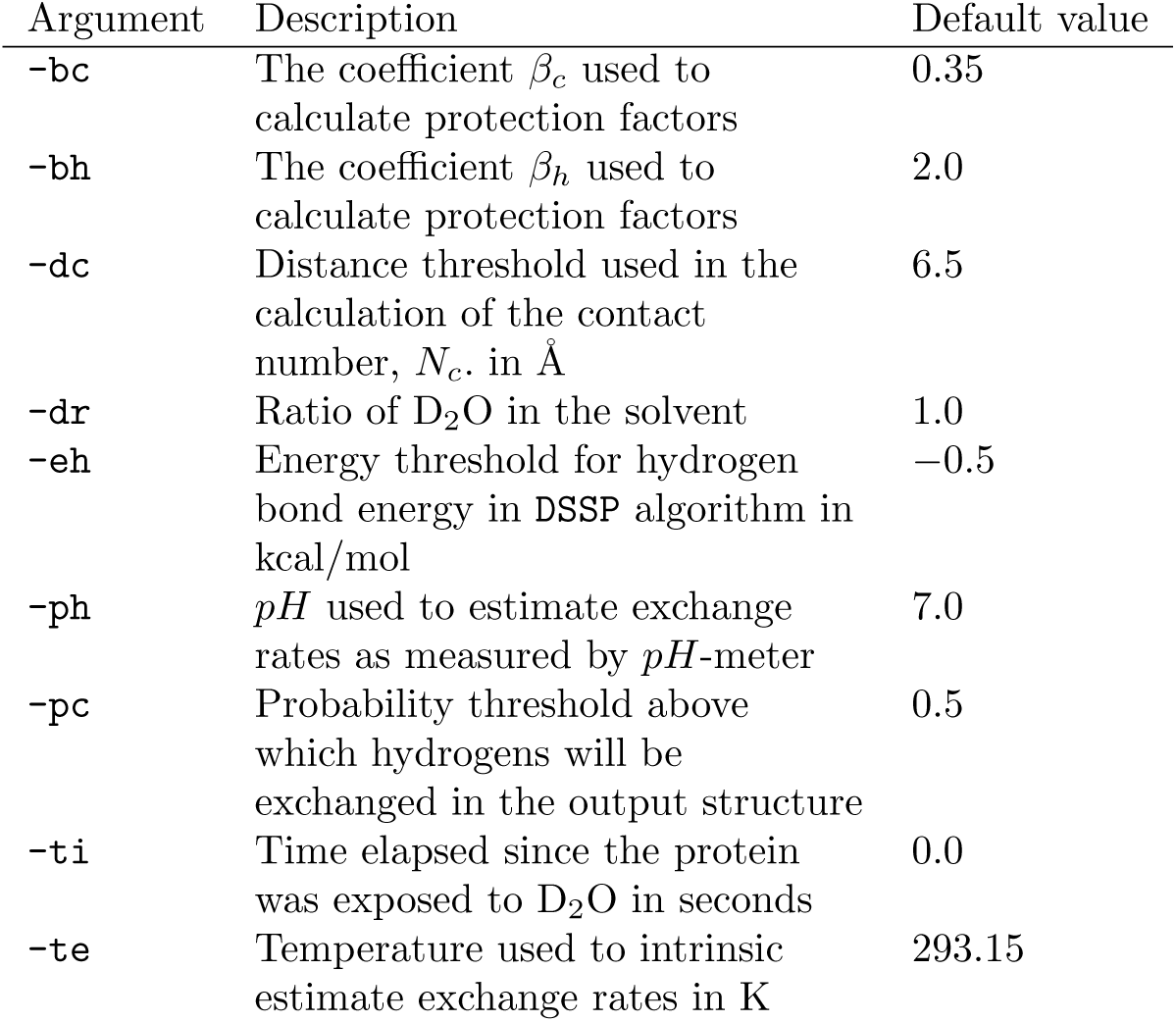
Selected optional command line arguments.

| Argument | Description | Default value |
| --- | --- | --- |
| -bc | The coefficient $\beta_c$ used to calculate protection factors | 0.35 |
| -bh | The coefficient $\beta_h$ used to calculate protection factors | 2.0 |
| -dc | Distance threshold used in the calculation of the contact number, $N_c$, in Å | 6.5 |
| -dr | Ratio of D<sub>2</sub>O in the solvent | 1.0 |
| -eh | Energy threshold for hydrogen bond energy in DSSP algorithm in kcal/mol | -0.5 |
| -ph | $pH$ used to estimate exchange rates as measured by $pH$-meter | 7.0 |
| -pc | Probability threshold above which hydrogens will be exchanged in the output structure | 0.5 |
| -ti | Time elapsed since the protein was exposed to D<sub>2</sub>O in seconds | 0.0 |
| -te | Temperature used to intrinsic estimate exchange rates in K | 293.15 |

The intrinsic exchange rates, *k*_int_, are calculated from the reference data published by the Englander Lab (Englander, 2019; Nguyen *et al*., 2018). As default, the software uses the reference values for poly-DL-Alanine; however, using their reference data from oligopeptides in the calculations is available as an alternative via a command line argument.

Whether hydrogens are present in the provided structure or not is irrelevant to the output. The software will remove any hydrogens from the structure and add them using the PDB2PQR software as well as the PROPKA algorithm.

The software outputs two files containing the provided structure. In one, values of *k*_int_ are exported as the B-factor column and values of log (*P_f_*) are exported as the occupancy column. The other file contains log (*k*_ex_) as the B-factors and *P*_ex_ as the occupancies. In the second structure, hydrogen atoms will have been exchanged to deuterium atoms if *P*_ex_ exceeds a preset threshold for the individual residues.

If several models are present in the given file (separated using the MODEL keyword), the protection factor calculation is done for each individual model and outputted. The protection factors are subsequently averaged as outlined in equation 1, and each model is deuterated according to this average in an effort to simplify using the software on e.g. ensembles from nuclear magnetic resonance or MD trajectories.

### 4.2. Code, dependencies, and availability

Source code for the presented software is available for download or cloning from this Gitlab repository: https://gitlab.com/mcpe/PSX

The software is published under version 3 of the GNU Publishing License, GPLv3, and is available ‘as is’.

The software is written in Python and relies on the DSSP software (Kabsch & Sander, 1983) for identifying hydrogen bonds and the PDB2PQR software for adding hydrogens to the structure based on the PROPKA algorithm. The reading and writing of the protein structures are handled using the PDB-tools (Hamelryck & Manderick, 2003) in the BioPython package (Cock *et al*., 2009). As such, these modules must be available for the software to run correctly.

In passing, we mention that for the examples shown in this paper the running time of the software is a couple of seconds on an ordinary laptop. The software has been tested on Ubuntu and OSX.

## 5. Perspectives and conclusions

We have presented software to estimate the effects of solvent exposure using well established models and assessed this effect in the context of small-angle neutron scattering. Future investigations will attempt to estimate these effects in the context of other scattering techniques; most notably neutron reflectometry, as this technique would be susceptible to and affected by inaccurate estimates of scattering length (densities). For neutron reflectometry in particular, H_2_O/D_2_O-contrast variations are done *in situ* and is usually an integrated part of the experiment. Thus, the outlined effects are especially relevant in this context.

Given recent developments and experiences in biological small-angle neutron scattering methods, we believe more precise approaches to the problem of solvent exchange are needed in the neutron scattering community in order to utilize the experimental set ups and the acquired data fully.

The model that we use to estimate protection factors could be optimized, including improved values of *β_c_* and *β_h_*. Refining optimal values for these parameters is an ongoing endeavour (Mohammadiarani *et al*., 2018) and future versions of the software might utilize different values, or indeed functional forms, than the one used here. If similar models are established for hydrogen-deuterium solvent exchange for e.g. RNA, DNA, or other relevant biomolecules the software could be generalized to include these systems as well.

To employ the outlined methods and ideas for analysis of neutron scattering data, we recommend that experimentalists record *pH*, temperature, and time exposed to D_2_O before and during the experiment. The calculations are primarily sensitive to the *pH* value, and secondarily to the time in which the sample has been exposed to D_2_O. From this information, using the outlined approach one will be able to accurately predict the deuteration levels in the sample.

Finally, we observe that the default assumption of 90% deuteration of proteins in SANS experiments appears to be a very high estimate for generic, modern BioSANS samples according to the considered model and typical temperatures and values of *pH* in experiments. Values between 50% and 70% seem more appropriate given our examples; however, ideally, these calculations should be repeated for each sample in an experiment. We hope the presented software will aid users in estimating this parameter.

## Appendix A Calculating scattering profiles

We use the Fast Debye sum approach to calculating scattering profiles begins by establishing a scattering-weighted pair distance distribution over *N* scatterers (Hansen, 1990):

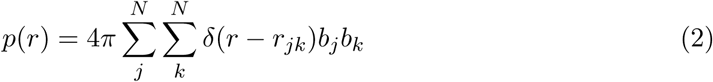

where *b_j_* and *b_k_* are the (excess) scattering lengths of the *j*’th and *k*’th scatterer, and *r_jk_*is the distance between them. In practice, this expression is usually binned into the data structure of a histogram. From this construction, we can calculate the scattering profile from the scatterers with a Fourier transformation:

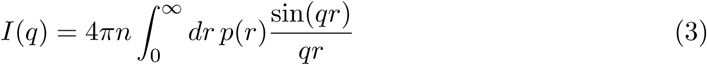

where *n* is the number density of the sample.

This method is readily extended for the structures emerging from the presented software.

If we (using e.g. the presented software) associate an exchange probability, *P*_ex_, to each backbone hydrogen (and set *P*_ex_ = 0 for all other scatterers), we get a concise expression by assigning two scattering lengths, *b*^H^ and *b*^D^, to each scatterer, where *b*^H^ is the scattering length in the case where the backbone hydrogen has not exchanged, and *b*^D^ is the scattering length elsewise. Note that *b*^H^ = *b*^D^ for the scatterers that are not backbone hydrogens.

If we assume that the *P*_ex_s are statistically independent, we can construct a substitute for equation 2:

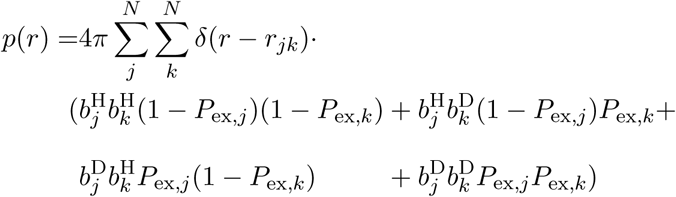

The Fourier transformation can now be carried out using equation 3.

## Appendix B Preparation and handling of BSA samples

### B.1. Sample preparation

BSA (Sigma Aldrich) was dissolved in gel filtration buffer (20 mM Tris-HCl, *pH* 7.5, 100 mM NaCl) and applied to a Superdex 200 Increase (GE Healthcare) equilibrated in gel filtration buffer. Fractions spanning the peak corresponding to monomeric BSA were pooled and kept on ice until exchange into D_2_O. The sample was split in four and each fraction changed into D_2_O using a NAP-5 desalting column (GE Healthcare).

The exchange into D_2_O was conducted by applying 0.5 ml sample to an equilibrated NAP5 column (20 mM Tris-DCl, 100 mM NaCl at either pH 7.3 or 8.8 in 100% D_2_O) and eluted with 0.95 ml buffer of the corresponding buffer. The initla drop was not collected, but the remaining eluate was collected and used for SANS measurements. The exchange was conducted either 9 or 36 hours before irradiation.

### B.2. Experimental conditions

The samples were irradiated by neutrons with a nominal wavelength of 6.0 Å from a distribution with a width of 0.6 Å in two experimental settings: one with a collimation length of 2.8 m and a sample-to-detector distance of 1.5 m, and one with a collimation length of 8.0 m with a sample-to-detector distance of 8.0 m covering a range in scattering momentum transfer from 0.0084 Å*^−^*^1^ to 0.57 Å*^−^*^1^. 1 mm rectangular Hellma Suprasil quartz cuvettes (Hellma Analytics) were usied as sample containers during the experiment.

Furthermore, absorption at 280 nm was measured on the irradiated samples using a Nanodrop 1000 spectrophotometer (Thermo Fisher Scientific), from which we estimate the protein concentration of the irradiated fraction. The extinction coefficient at 280 nm for a BSA monomer was calculated to be 42925 M*^−^*^1^cm*^−^*^1^.

Instrumental effect were accounted for as described in the literature (Pedersen *et al*., 1990) using the values produced by GRASP (Dewhurst, 2019), which was also used to reduce the data and bring them to absolute units using the direct beam chopped by an ultra-fast chopper as reference. The presented models were calculated and instrumental effects were taken into account using WillItFit (Pedersen *et al*., 2013).

We thank the staff at Institut Laue-Langevin, Grenoble, France for the beam time and support resulting in the presented data, and thank Nicolai Tidemand Johansen for aiding in the preparation and irradiation of the BSA sample. We thank Raul Araya-Secchi for discussions on general implementations of the selected algorithms. This research was supported by the Lundbeck Foundation BRAINSTRUC initiative in structural biology, and the Synergy project funded by the Novo Nordisk foundation. NVidia’s Academic Grant Program is acknowledged for granting the Titan Xp GPU on which many of the presented calculations were performed.

**Synopsis** Using established models, we calculate the effect of exposing protonated protein samples to D_2_O and assess the impact in small-angle neutron scattering experiments. We introduce software to evaluate the effect of this from a given structure and give examples of applications.

